# Photopatterned Sacrificial Vascular Architectures for Large Tissue-Scale Oxygenation

**DOI:** 10.64898/2026.02.27.708614

**Authors:** Ian A. Coates, Carl A. Kohnke, Yee Lin Tan, Daniel I. Alnasir, Annie Nguyen, Elbert E. Heng, Austin Kwan, Maria T. Dulay, Bruce Schaar, Mark A. Skylar-Scott, John W. MacArthur, Eric S.G. Shaqfeh, Joseph M. DeSimone

## Abstract

The engineering of thick, metabolically active tissues is constrained by the lack of scalable methods to create perfusable vasculature. This hinders effective metabolite transport in large tissue volumes, posing a critical barrier for regenerative tissue applications. In this study, we introduce photopatterned Channel Architectures with Sacrificial Templates (pCAST), an additive manufacturing strategy for generating three dimensional (3D), interconnected vascular networks with precisely defined negative space. Water-soluble sacrificial templates were fabricated using scalable Continuous Liquid Interface Production (CLIP), embedded within tissue constructs, and flushed away to yield 50 µm perfusable channels spanning centimeter-scale tissue constructs. We then apply experimental oxygen mapping and viability analysis to pCAST constructs to build finite-element models that predict patterns of oxygen availability and tissue survival are governed by the balance between metabolic demand and vascular architecture, consistent with reaction–diffusion theory. This computational framework quantitatively predicts oxygen distributions and viability boundaries across vascular geometries and is validated experimentally. Together, these results establish pCAST as a scalable design framework linking vascular architecture, perfusion, and metabolic support for engineering large, 3D perfused tissue constructs.

**Significance:** The ability to engineer thick, living tissues is limited by poor oxygen and nutrient delivery, which causes cell death before tissues can function or integrate with the body. This work addresses that fundamental barrier by introducing photopatterned Channel Architecture with Sacrificial Templates (pCAST), a scalable manufacturing strategy that creates precisely defined, perfusable vascular networks inside 3D tissues. By combining high-resolution 3D printing, sacrificial templating, and quantitative oxygen mapping, this research establishes design rules that link vascular geometry, perfusion, and tissue viability. These insights provide a general framework for building large, metabolically active tissues, with direct relevance to cardiac patches and other regenerative medicine applications.

## Introduction

The fabrication of large, functional tissues remains fundamentally constrained by mass transport (*1*–*3*). Oxygen must be continuously replenished to support cellular metabolism yet diffuses only a few hundred micrometers in dense cellular tissues (*4*–*6*). Engineered tissue constructs that lack integrated vasculature therefore develop steep oxygen gradients, leaving only a thin rim of viable cells surrounding perfused regions (*7*–*11*). Native tissues overcome this limit through hierarchical vascular networks that couple convective transport in vessels to diffusive exchange in surrounding parenchyma (*12*–*14*). Reproducing this multiscale organization is essential for sustaining physiologic cell densities in engineered tissues and for advancing applications in regenerative medicine, disease modeling, and drug discovery (*2, 15*–*18*).

To overcome transport limitations, native tissues organize blood supply through hierarchical vascular architectures in which a single artery–vein pair supplies and drains a well-defined three dimensional (3D) tissue volume, forming an angiosome in which oxygen delivery is matched to local metabolic demand (*19, 20*). This organization – where a few large vessels feed many smaller branches – provides an efficient engineering strategy for sustaining large tissue volumes while maintaining simple, tractable boundary conditions. Recent computational studies have leveraged this principle to design synthetic vascular networks by prescribing branched geometries and modeling oxygen transport within digitally defined vessels (*8, 21, 22*). However, these efforts have largely emphasized intravascular transport, with comparatively limited treatment of oxygen diffusion and consumption throughout the surrounding tissue volume, which ultimately governs viability. It is our contention that engineering viable large-scale tissues therefore will require vascular architectures that couple single-inlet, single-outlet perfusion with distributed branching capable of sustaining tissue-level oxygen transport across the full construct.

Reproducing this hierarchical organization in engineered tissues has motivated a range of fabrication strategies enabled by additive manufacturing (AM), which allows 3D architectures to be specified with spatial precision reflecting vascular design principles. Early templating approaches such as soft lithography and needle molding established the feasibility of forming defined negative spaces in hydrogels, though their geometries were restricted (*23*–*26*). Extrusion-based bioprinting increases design flexibility but faces trade-offs between resolution and throughput (*7, 21, 27*–*29*). Direct photopolymerization fabrication of channel lumens, including digital light processing (DLP) and two-photon polymerization (2PP), achieve high spatial fidelity but are limited to photocurable matrices that limit cell compatibility and remodelability and are difficult to scale to large, cell-dense constructs (*30, 31*). Sacrificial templating, in contrast, offers broader material compatibility, as a broad range of biological or synthetic gels may be casted around a printed fugitive structure, which upon removal yields perfusable networks. However, the fidelity and scalability of photopatterned fugitive networks remains limited (*8, 29, 32, 33*).

To meet this challenge, we introduce pCAST (photopatterned Channel Architectures with Sacrificial Templates), a sacrificial templating platform in which vascular architectures are fabricated using Continuous Liquid Interface Production (CLIP), a vat photopolymerization AM approach, to achieve scalable micron-scale resolution throughout volumes with dimensions on the centimeter-scale to support the fabrication and sustained viability of tissue constructs (*34, 35*). The pCAST approach starts with generating the vascular-like network made from a dissolvable template, which is then encapsulated in a cell matrix medium. Subsequently, the vascular template is rapidly dissolved (typically within 5 minutes) under mild, cell-compatible conditions to yield perfusable vascular networks that can distribute oxygen within minutes after cellular matrix deposition.

To rigorously assess the quality and performance of our synthetic angiosome-like constructs fabricated by pCAST, we developed an experimentally validated finite-element model (FEM) simulation framework to resolve time-dependent, three-dimensional oxygen transport within these vascularized tissue constructs. Using this model in combination with real-time oxygen imaging and confocal microscopy, we systematically examined how metabolic demand, vascular surface-area density, and network topology govern tissue-scale oxygen delivery and cellular perfusion in large, perfused constructs. We show that pCAST-created constructs bridge computational vascular design and experimental tissue engineering, providing a practical framework for engineering angiosome-scale perfusion which could enable future applications in in vitro disease modeling and regenerative tissue grafting.

## Results

### Fabrication of Sacrificial Templates for High-Resolution 3D Vascular Architectures

To enable the fabrication of sacrificial vascular templates with complex 3D topology, we developed a water-soluble sacrificial resin designed to simultaneously satisfy the competing requirements of high-resolution photopolymerization, mechanical robustness, biocompatibility, and rapid dissolution following template encapsulation in a hydrogel matrix. Balancing these constraints is critical for producing freestanding vascular templates that can be transferred intact into surrounding matrices and subsequently sacrificed to yield perfusable negative-space networks.

The sacrificial resin formulation was 4-acryloylmorpholine (ACMO), a hydrophilic monofunctional acrylamide selected for its low viscosity, rapid aqueous solubility, biocompatibility, and compatibility with light based fabrication processes (*36*). Phenylbis(2,4,6-trimethylbenzoyl)phosphine oxide (BAPO) was used as the photoinitiator, and the formulation was tuned to balance print fidelity with biocompatibility (Fig. S5) and mechanical integrity. Following aqueous perfusion, the polymerized ACMO templates dissolved completely within minutes, enabling rapid and on-demand removal without disrupting the surrounding construct (Fig. 1).

**Fig. 1.**
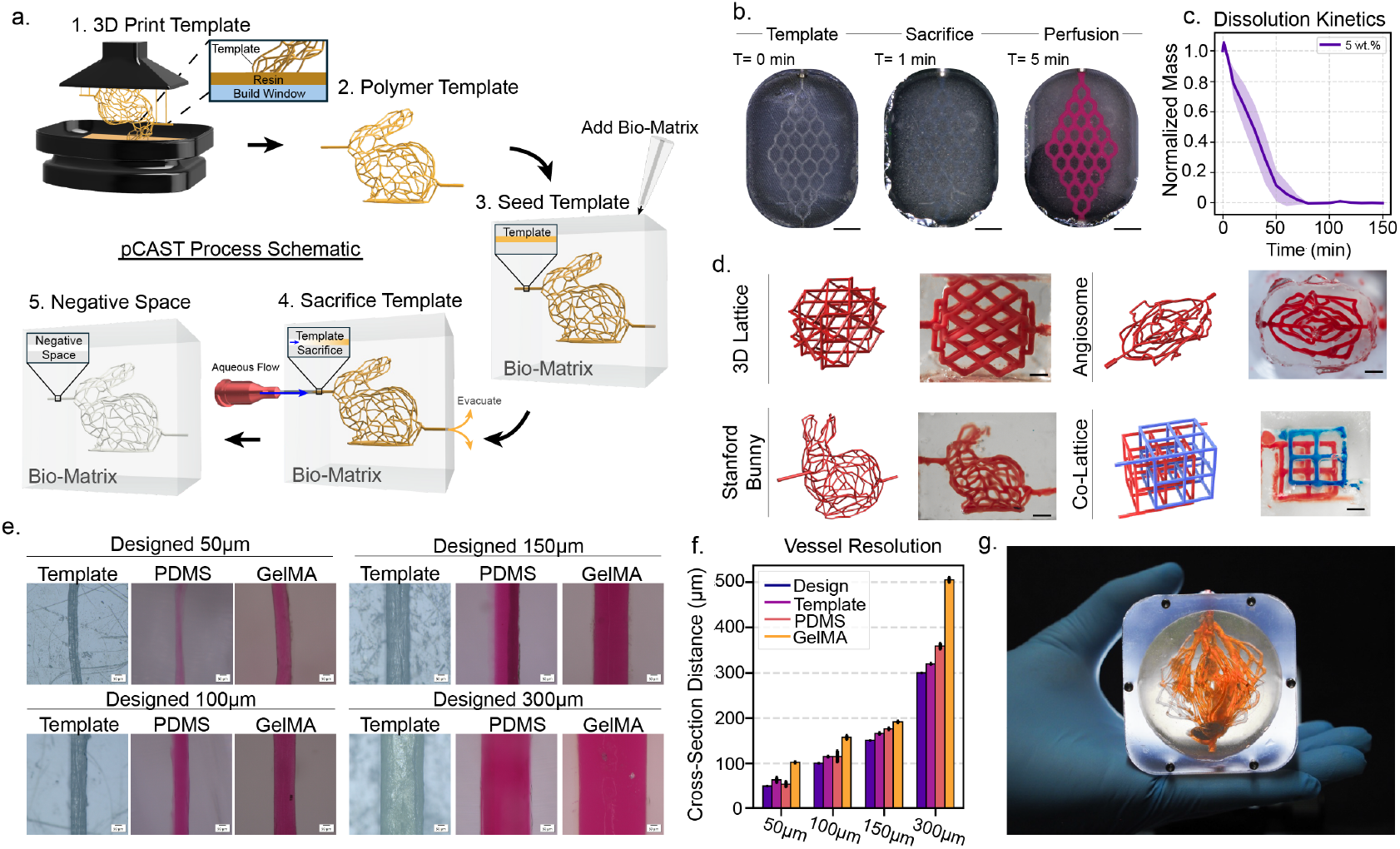
Scalable fabrication of perfusable vascular architectures using the pCAST process. **(a)** Schematic of the pCAST workflow. (**b)** Time-lapse demonstration of template embedding, sacrifice, and media perfusion within a hydrogel construct. Scale bars: 2.5 mm. **(c)** Dissolution kinetics of ACMO-based sacrificial resin. Mean ± SD (N=3). **(d)** 3D printed vascular templates and corresponding negative-space networks after sacrifice. Scale bars: 1 mm. **(e)** Fidelity of channel dimensions across PDMS molds and GelMA hydrogels for designed features ranging from 50–300 µm. Scale bars: 50 µm. **(f)** Vessel resolution quantification. Mean ± SD (N=3). **(g)** Large-volume perfused construct housed in a bioreactor.

Sacrificial templates were printed directly from computer-aided design (CAD) geometries using a 5.4 µm pixel CLIP AM process. The oxygen-inhibited “dead zone” intrinsic to CLIP enables scalable high-resolution template fabrication across length scales (*34, 35*). Printed templates are embedded within hydrogel matrices and subsequently sacrificed via aqueous perfusion to yield interconnected negative-space channel networks (Fig. 1a,b). pCAST successfully scaled across complex, multi-bifurcating architectures with interpenetrating dual networks, from sub-centimeter constructs (<0.5 cm^3^) to large constructs (>36 cm^3^), while maintaining geometric fidelity and robust perfusion (Fig. 1d,g).

Next, we evaluated print resolution across designed channel diameters ranging from 50 to 300 µm. Template features as small as 50 µm lumen diameters were reproducibly fabricated and successfully transferred into both PDMS molds and GelMA hydrogels (Figs. 1e,f). Across the final lumen diameters spanning 50–300 µm, with mean deviations from scaffold template of 15% (PDMS) and 64% (GelMA). Deviation is likely due to swelling of the polymeric sacrificial templates during dissolution leading to increased volume. Additional factors, including the stiffness and hydration state of the surrounding matrix influence the degree of enlargement of the resulting hollow lumens.

### Finite-Element Modeling of Perfusion Oxygen Transport in pCAST Constructs

Because oxygen is a rapidly consumed, non-storable metabolic substrate with low solubility in aqueous media, its delivery imposes a dominant transport constraint that ultimately limits metabolic support and cell viability in engineered tissues (*9, 37*–*39*). We therefore focus on oxygen as a primary design variable for evaluating vascular performance within pCAST constructs. To quantify oxygen delivery within pCAST constructs, we developed a physics-based finite-element model (FEM) describing oxygen exchange between perfused pCAST vascular channels and the surrounding engineered tissue. Convective transport within channel lumens is coupled to diffusive transport and cellular consumption in the hydrogel domain, enabling quantitative prediction of dissolved oxygen concentration throughout the tissue volume, not only within the vasculature (Eq. 1). Cellular oxygen uptake is represented as a distributed sink term within the gel and modeled using Michaelis–Menten kinetics (Eq. 1) (*40*–*42*).

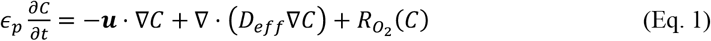

In Eq.1, *C*(*x, t*) is dissolved oxygen concentration; ϵ_*p*_ is the hydrogel porosity; *u*(*x, t*) is the local superficial velocity obtained from the flow solution; *D*_*eff*_ is the effective oxygen diffusivity accounting for porosity/tortuosity; and 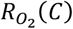 is the local reaction term, taken as zero in acellular gels and negative for cellular consumption. Concentration and partial pressure are related through Henry’s Law, 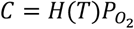. Details of the Henry coefficient, diffusivity scaling, governing flow equations, and boundary conditions are provided in Table S1.

This modeling framework provides a mechanistic basis for interpreting experimentally measured oxygen gradients within tissue constructs, enabling direct, quantitative comparison between predicted and observed oxygen distributions across vascular architectures and cell densities. Importantly, the framework resolves oxygen transport and consumption throughout the surrounding tissue matrix rather than restricting analysis to intravascular oxygen levels, capturing the spatial gradients that govern bulk cellular viability.

### Real-Time Mapping of Oxygen Gradients in Acellular pCAST Networks

To isolate the transport contribution of convection and diffusion, we first characterized oxygen delivery in acellular pCAST constructs where 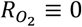. Constructs containing defined vascular channels were embedded in 10 wt.% GelMA and mounted in a sealed perfusion chamber equipped with an oxygen-sensitive luminescent sensor foil positioned beneath the gel (Fig. 2a). The detector records spatiotemporal oxygen distributions via luminescence quenching, wherein increasing dissolved oxygen reduces emission intensity in a calibrated manner. This approach enables non-invasive, real-time mapping of oxygen concentration within the tissue rather than being limited to intravascular measurements.

**Fig. 2.**
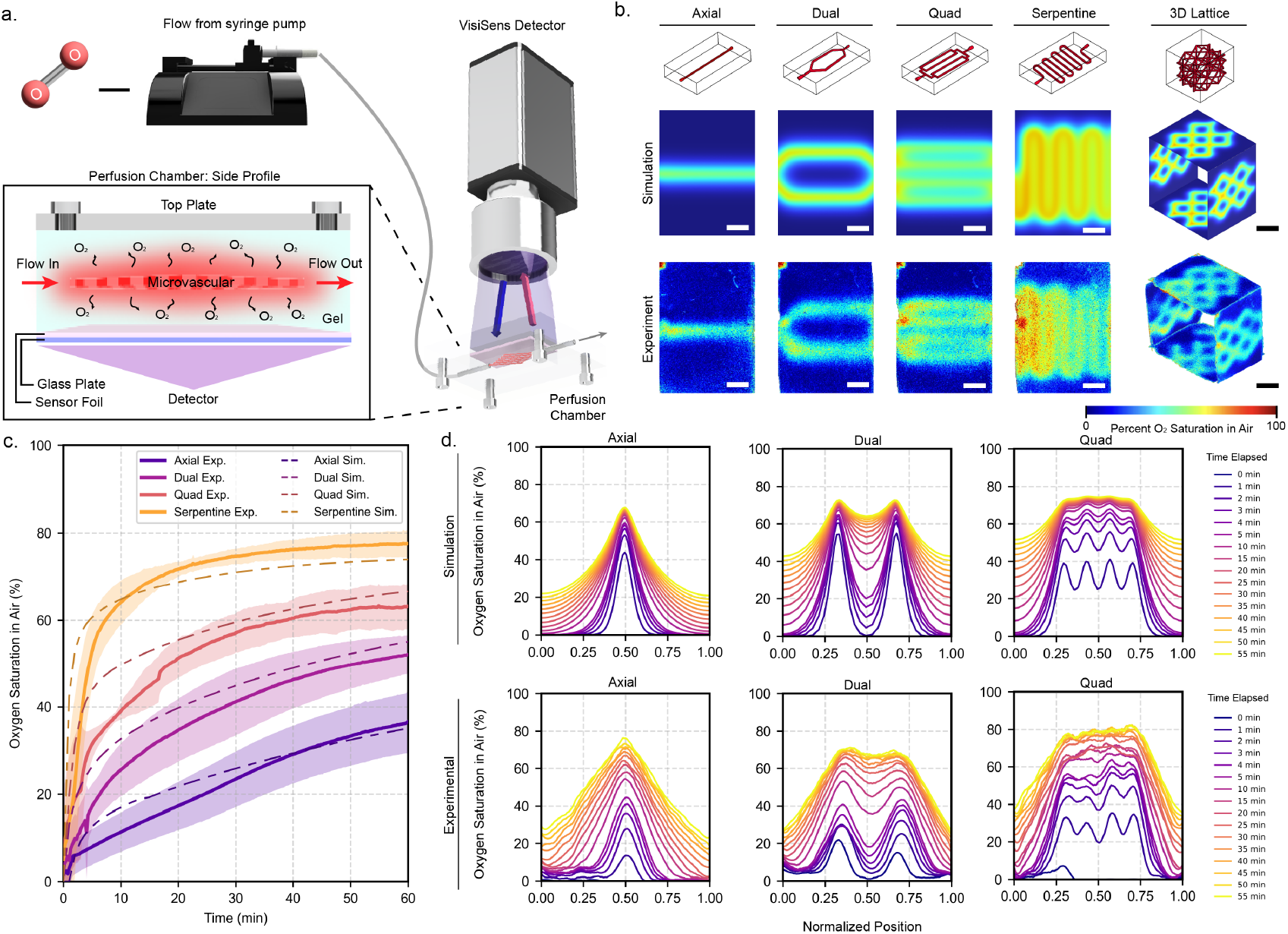
Integration of perfusion imaging with finite-element oxygen transport modeling in pCAST microvascular networks. **(a)** Schematic of the perfusion platform used for oxygen imaging. (**b)** Comparison of simulated and experimentally measured oxygen fields in acellular constructs. Scale bars 1 mm. **(c)** Temporal oxygen saturation measured experimentally (solid traces with shaded SD) and predicted by FEM (dashed) for each architecture. Mean ± SD (N=3). **(d)** Cross-sectional oxygen profiles over time for axial, dual, and quad geometries in simulation (top) and experiment (bottom). Mean ± SD (N=3).

We evaluated four representative vascular geometries, axial, dual, quad, and serpentine, which span increasing vascular network density and perfusion coverage (Fig. 2b). Time-resolved oxygen maps revealed that oxygen enters the matrix from the channel lumens and spreads outward into the hydrogel domain. Architectures with denser networks (dual, quad, serpentine) exhibited more rapid and spatially uniform oxygenation, whereas the single axial channel generated steeper gradients extending into the bulk.

Quantitative traces of mean oxygen saturation over time demonstrated that networks with greater surface-area density reached higher steady-state oxygen levels more quickly (Fig. 2c). Cross-sectional oxygen profiles orthogonal to the channels showed that increasing the number of channels reduced gradient steepness and expanded the region of high oxygen availability within the matrix (Fig. 2d), indicating more uniform tissue-level oxygen delivery.

FEM simulations based on Eq. 1 and Eqs. S1-S4 were constructed using the measured channel geometry, inlet oxygen concentration, flow rate, and hydrogel diffusion coefficient. The FEM model reproduced both temporal oxygen rise and spatial cross-sections (dashed curves, Figs. 2c,d), confirming that our FEM simulations and experiments are mutually cross-validated and accurately describe transport behavior in acellular networks.

### Predicting and Measuring Oxygen Gradients in Living pCAST Tissues

We next examined cell-laden pCAST constructs 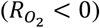 to determine how metabolic demand interacts with vascular architecture to regulate oxygen availability in engineered tissues. HEK-293T cells were encapsulated in 10 wt.% GelMA at densities of 15, 30, 60, and 100 million cells mL^−1^, and constructs were perfused with Dulbecco’s Modified Eagle’s Medium (DMEM) based cell media at a constant flow rate of 250 µL min^−1^ (Fig. 3a). Dissolved oxygen fields were monitored continuously using planar luminescence-quenching sensors, enabling real-time mapping of oxygen gradients within living hydrogels and reported after 24h (Figs. 3b,c).

**Fig. 3.**
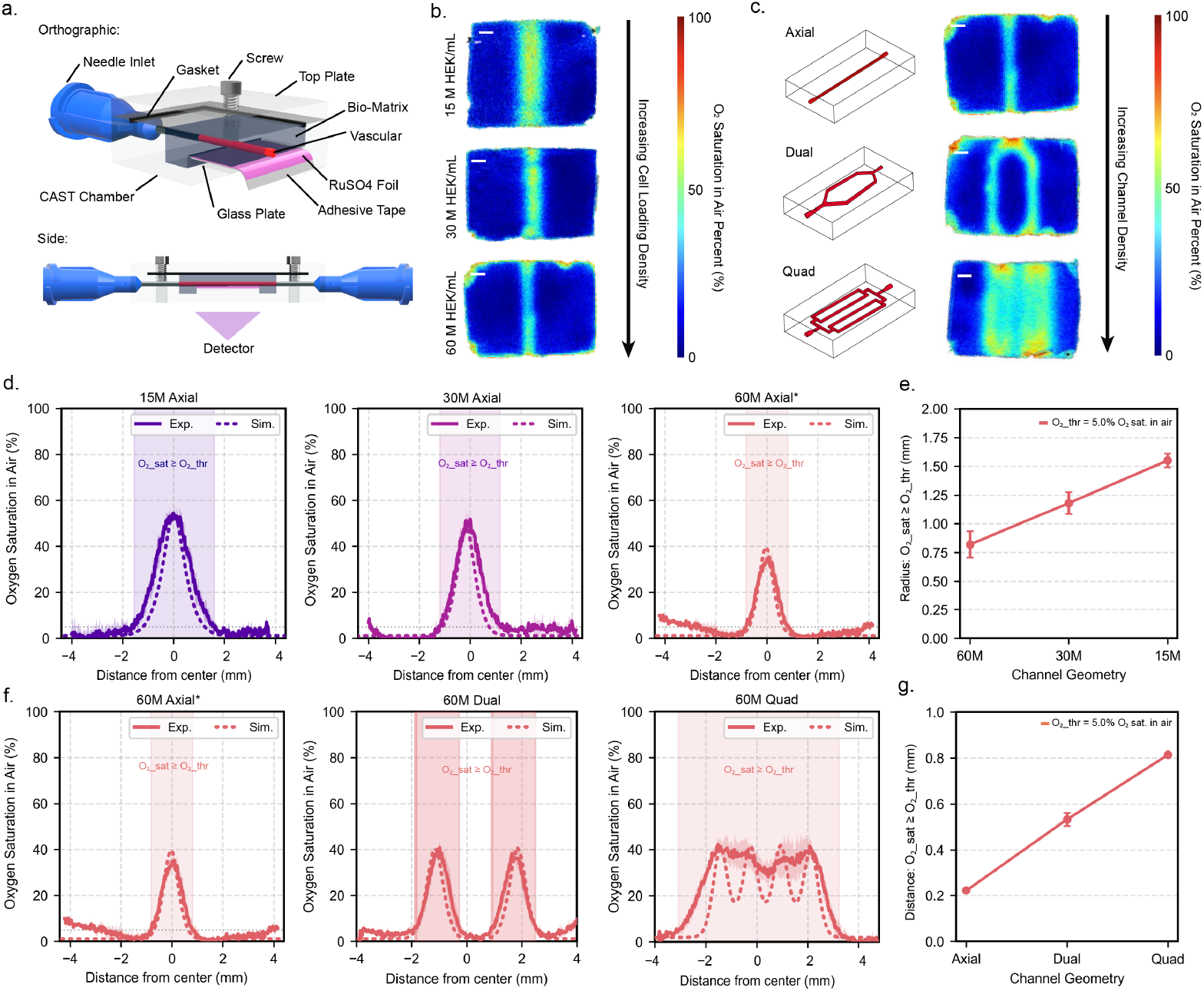
Oxygen transport in perfused pCAST constructs reveals coupled effects of cell density and vascular geometry on oxygen penetration. **(a)** Schematic of the pCAST perfusion chamber configured for real-time oxygen mapping using an oxygen-sensitive luminescent sensor. **(b)** Experimentally measured cross-sectional oxygen saturation maps for axial-channel constructs containing increasing HEK-293T cell densities (15, 30, and 60 million cells mL^−1^). Scale bars 1 mm. **(c)** Comparison of axial, dual, and quad channel architectures at a fixed cell density of 60 million cells mL^−1^. Scale bars 1 mm. **(d)** Radial oxygen saturation profiles for axial constructs across increasing cell densities, shown alongside corresponding FEM simulations. Shaded regions indicate distances over which oxygen saturation remains above a one percent oxygen threshold. Mean ± SD (N=3). **(e)** Quantification of the oxygen penetration distance, defined as the radial distance from the channel wall over which oxygen saturation remains above the one percent threshold, as a function of cell density. Mean ± SD (N=3). **(f)** Radial oxygen saturation profiles for axial, dual, and quad channel architectures at 60 million cells mL^−1^, comparing experimental measurements and FEM predictions. Mean ± SD (N=1). **(g)** Oxygen penetration distance as a function of vascular geometry evaluated across multiple oxygen saturation thresholds. Mean ± SD (N=1).

At a fixed axial channel geometry, increasing cell density progressively narrowed the oxygenated region surrounding the perfused lumen and steepened oxygen gradients into the surrounding matrix (Fig. 3b,d). The oxygen penetration radius, defined as the distance from the channel edge over which oxygen saturation remained above a 1 percent oxygen threshold, decreased monotonically with increasing cell density (Fig. 3e) (*39*). This behavior is consistent with reaction–diffusion scaling, where the characteristic oxygen penetration length 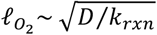decreases as volumetric cellular consumption increases. Under fixed perfusion conditions, metabolic demand therefore becomes the dominant factor that governs oxygen penetration depth.

Increasing vascular network density partially mitigated this limitation: dual and quad channel architectures produced broader and more uniform oxygen distributions than axial networks at matched cell densities (Fig. 3c,f) by reducing mean diffusion distances and effectively extending the reaction–diffusion length scale, resulting in an expanded oxygen-sufficient region with increasing channel density (Fig. 3g).

FEM modeling incorporating Michaelis–Menten cellular consumption quantitatively reproduced both absolute oxygen levels and spatial oxygen profiles across cell densities and vascular geometries (Fig. 3d,f), demonstrating that oxygen availability in pCAST constructs arises from the balance between cellular metabolic demand and geometry-controlled diffusive transport. Extension of this framework to additional parameters, including cell type–dependent consumption rates, matrix porosity, and perfusion flow rate (Fig. S3) further establishes its general applicability across biologically relevant conditions.

### Cell Viability Reflects the Spatial Structure of Oxygen Delivery in pCAST Tissues

To directly link oxygen transport to cell survival in the bulk matrix, we quantified spatial cell viability in cell-laden pCAST constructs following 5 days of extended perfusion culture (Fig. 4a).

**Fig. 4.**
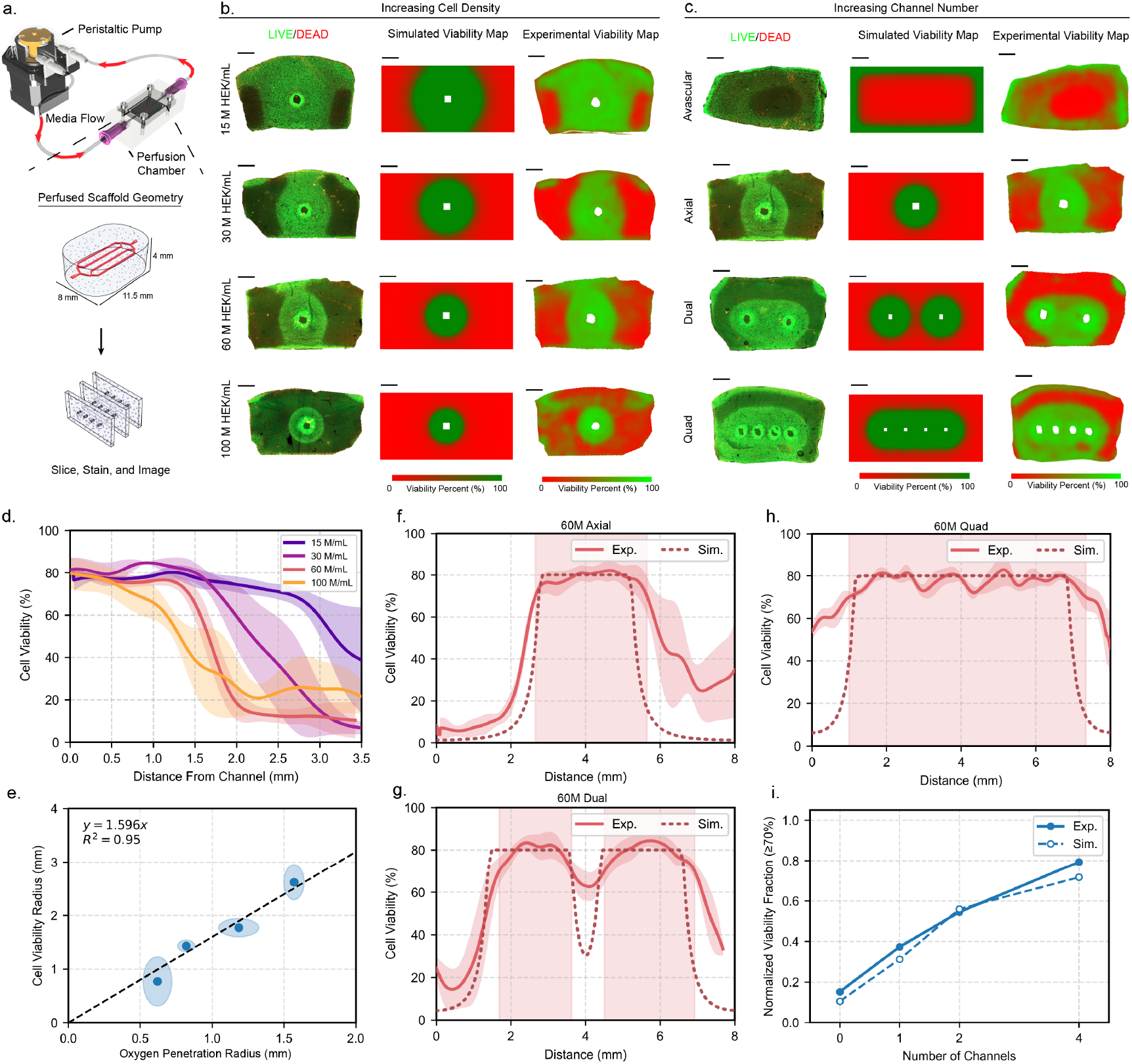
Oxygen-limited cell viability in perfused pCAST constructs is governed by cell density and vascular architecture. **(a)** Schematic of the perfusion platform used for long-term culture, consisting of a peristaltic pump, perfusion chamber, and pCAST-fabricated vascular templates embedded in GelMA hydrogels, followed by sectioning and LIVE/DEAD staining after culture. **(b)** Effect of increasing cell density (15–100 million HEK-293T cells mL^−1^) in constructs containing a single axial channel. Shown are representative LIVE/DEAD fluorescence images (left), FEM-predicted viability maps assuming hypoxic cell death below a one percent oxygen threshold (center), and experimentally measured viability maps (right). Scale bars 1 mm. **(c)** Effect of increasing channel number at fixed cell density (60 million cells mL^−1^). Representative LIVE/DEAD fluorescence images (left), FEM-predicted viability maps assuming hypoxic cell death below a one percent oxygen threshold (center), and experimentally measured viability maps (right). Scale bars 1 mm. **(d)** Radial viability profiles for axial constructs across increasing cell densities. Mean ± SD (N=3). **(e)** Correlation between the experimentally measured cell viability radius and the oxygen penetration distance derived from oxygen mapping. Mean ± SD (N=3). **(f–g)** Comparison of experimental and FEM-predicted radial viability profiles for axial, dual, and quad vascular geometries at 60 million cells mL^−1^. Shaded regions indicate distances over which modeled oxygen concentrations remain above the hypoxic death threshold. Mean ± SD (N=3). **(i)** Experimentally measured and FEM-predicted normalized viable tissue fraction for cellular templates at 60 million cells mL^−1^.

HEK-293T cells were encapsulated in 10 wt.% GelMA at densities of 15, 30, 60, and 100 million cells mL^−1^ and perfused continuously with cell media at 250 µL min^−1^ for 5 days using a closed-loop perfusion system. After culture, constructs were sectioned and stained with LIVE/DEAD (Calcein-AM/EthD-1) dyes to generate spatially resolved viability maps and radial viability profiles relative to perfused channels (Fig. 4b-d).

Given a fixed axial geometry, increasing cell density produced a progressive contraction of the viable tissue region surrounding the channel (Fig. 4b). At high cell densities reflective of native tissues (~100 million cells mL^−1^), viability dropped sharply beyond a narrow perivascular zone, consistent with hypoxia-driven loss of viability. Radial viability profiles revealed steep transitions from high to low viability that shifted closer to the channel with increasing density, indicating that metabolic demand increasingly outpaced oxygen delivery (Fig. 4d). In this discussion, we define the viability radius as the radial distance at which the measured cell viability decreased by more than 10 percent relative to perivascular values, forming a distinct “viability ring” around the channel.

To quantitatively relate oxygen availability to cell viability, oxygen penetration radius previously determined in the pCAST constructs were compared with the viability radius from matched geometries. Both the oxygen penetration distance and the viability radius decreased monotonically with increasing cell density and exhibited a strong, near-linear correspondence (Fig. 4e), suggesting that oxygen availability functions as the dominant limiting reagent governing spatial viability patterns under these perfusion conditions. Deviations from a strict one-to-one relationship likely reflect additional metabolic stressors not captured in the oxygen-only model, including nutrient depletion, waste accumulation, and heterogeneous cellular states.

We next evaluated whether vascular architecture could mitigate density-dependent viability loss. At a fixed density of 60 million cells mL^−1^, constructs incorporating dual and quad channel networks exhibited substantially broader and more uniform viability profiles compared to axial geometries (Figs. 4c,f–h). Increasing channel density reduces mean diffusion distances from the channels to cells, thereby expanding the region over which oxygen remained above the hypoxic threshold. Using the FEM simulations, binary spatial cellular viability (live or dead) was predicted directly from simulated oxygen fields by applying a defined hypoxic cutoff, classifying regions as viable or non-viable based on whether local oxygen concentrations exceeded the 1 percent threshold (Figs. 4b,c). The resulting FEM predicted viability profiles closely matched both the shape and spatial extent of the experimentally measured LIVE/DEAD profiles across all vascular geometries (Figs. 4f–h), supporting that oxygen transport and cellular oxygen consumption are primary contributors to viability patterns in these systems.

Collectively, these results demonstrate that cell survival in pCAST constructs is governed by a predictable balance between metabolic demand and geometry-controlled oxygen delivery. By explicitly linking oxygen penetration distance to independently measured viability loss over multi-day perfusion, this framework establishes quantitative design rules for vascular architectures capable of sustaining dense, clinically relevant tissue constructs.

### Computational Design, Fabrication, and Perfusion of Biomimetic Vascular Networks

Building on the ability to fabricate complex, perfusable vascular architectures with pCAST and to quantitatively predict tissue-scale oxygen distributions using an experimentally validated FEM simulation, we next sought to design 3D vascular networks that explicitly satisfy metabolic demands throughout large tissue volumes. To this end, we implemented a variant of the mutual tree attraction (space colonization) algorithm, previously applied to venation modeling and vascular morphogenesis, to generate hierarchical vascular architectures under defined manufacturability and perfusion constraints (*43, 44*). In this framework, vessel trees initiated from prescribed inlet and outlet seeds iteratively extend toward randomly distributed attraction points; while competing growth fronts exert mutual influence that promotes convergence into a continuous, space-filling dendritic network. This process yields hierarchical vascular geometries with locally adaptive branching and distributed path lengths predicted by FEM to keep entire tissue volume viable (Fig. 5a).

**Fig. 5.**
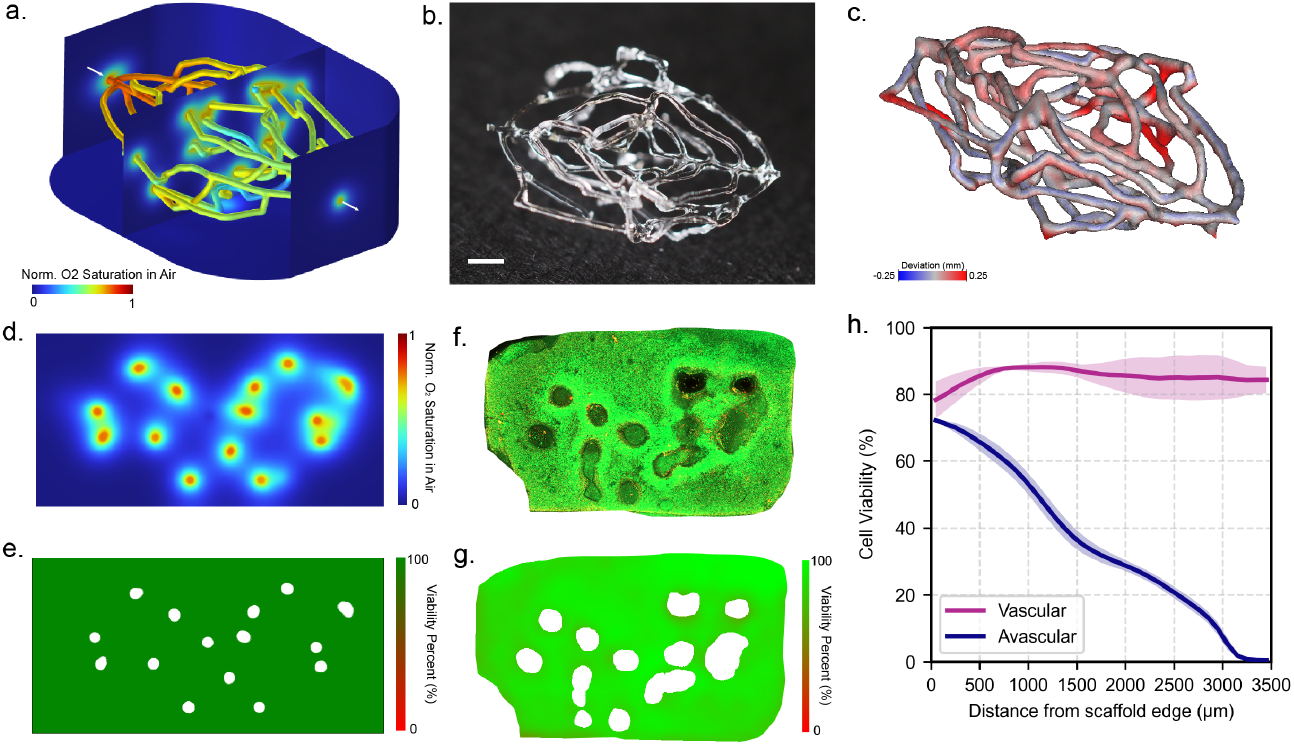
Computational design, fabrication, and functional validation of large-scale pCAST vascular templates. **(a)** Finite-element simulation of oxygen transport profiles within a volumetric pCAST generated construct containing a biomimetic vascular network. **(b)** Photograph of a CLIP-printed sacrificial vascular template generated using a space-colonization–based growth algorithm. Scale bar 1.5 mm. **(c)** Surface deviation map comparing the printed sacrificial template to the corresponding digital model, demonstrating high geometric fidelity across the construct volume. **(d)** Representative simulated two-dimensional slice of the oxygen saturation field extracted from the volumetric model. **(e)** Representative simulated two-dimensional slice of the predicted cell viability field based on oxygen-dependent consumption and death thresholds. **(f)** Representative LIVE/DEAD fluorescence image of a large, perfused, cell-laden pCAST construct following multi-day culture. **(g)** Quantitative spatial viability heat map derived from fluorescence image segmentation. **(h)** Radial viability profiles comparing vascularized and avascular constructs. Mean ± SD (N=3).

The resulting designs were exported directly to the pCAST workflow and fabricated as free-standing sacrificial templates (Fig. 5b). 3D reconstruction and deviation mapping confirmed close agreement between the computational design and printed structures, with sub–hundreds-of-microns deviations across the network volume (Fig. 5c).

Embedding these vascular templates within cell-laden hydrogel constructs enabled direct experimental validation of the model-guided designs. FEM simulations predicted spatial oxygen saturation fields arising from the complex, branched channel topology, revealing overlapping diffusion zones and distributed oxygen sources throughout the construct cross-section (Fig. 5d). Corresponding experimental oxygen mapping and LIVE/DEAD viability imaging closely matched these computational predictions (Figs. 5f,g). Quantitative viability analysis demonstrated substantially improved survival in vascularized constructs compared to avascular controls, which exhibited progressive loss of viability with increasing distance from the construct boundary (Fig. 5h). Together, these results demonstrate that algorithmically generated vascular networks, designed using a space colonization framework and evaluated using a 3D, experimentally validated oxygen transport model, can be reliably manufactured and function as distributed oxygen delivery systems capable of sustaining metabolically demanding 3D engineered tissues.

## Discussion

Oxygen transport remains a fundamental constraint in engineering thick, metabolically active tissues. In this study, we establish pCAST as a scalable fabrication platform capable of producing complex, tissue-scale vascular architectures with microscale fidelity and robust perfusion compatibility. By integrating this fabrication capability with real-time oxygen mapping and an experimentally validated FEM simulation framework, we provide a predictive approach for relating vascular geometry and cellular demand to sustaining oxygen availability.

The dominant role of oxygen in constraining tissue viability is well established across engineered constructs and native myocardium. Prior experimental and modeling work demonstrate that oxygen transport, rather than glucose or other soluble nutrients, defines the characteristic diffusion-limited penetration depth in metabolically dense tissues, primarily due to its limited solubility relative to cellular demand rather than diffusivity alone (*9, 37–39*). Although glucose diffuses more slowly than oxygen, it is not typically rate-limiting because its critical concentration and consumption rate remain low relative to its available concentration. Consistent with prior studies, oxygen therefore imposes a characteristic penetration length that directly constrains viable tissue volume; here, we extend this framework by directly mapping spatial oxygen gradients and correlating them with viability within architected vascular trees, establishing capillary geometry as a transport-specified design parameter rather than an empirical choice.

Extending vascularized constructs to larger volumes is critical not only for reducing edge-dominated culture artifacts, but also for enabling tissue dimensions relevant to graft integration, organ-scale disease models, and, ultimately, vascularized tissue replacement strategies where perfusion must be sustained immediately upon implantation. Building on these insights, we demonstrate a preliminary extension of pCAST to a substantially larger (~36 cm^3^), biomimetic vascular construct generated using an algorithmic tree-based design approach (Fig. 6). The resulting network supported perfusion and high average viability across millimeter-scale distances from the template edge, establishing feasibility for scaling pCAST-guided designs beyond simple channel layouts.

**Fig. 6.**
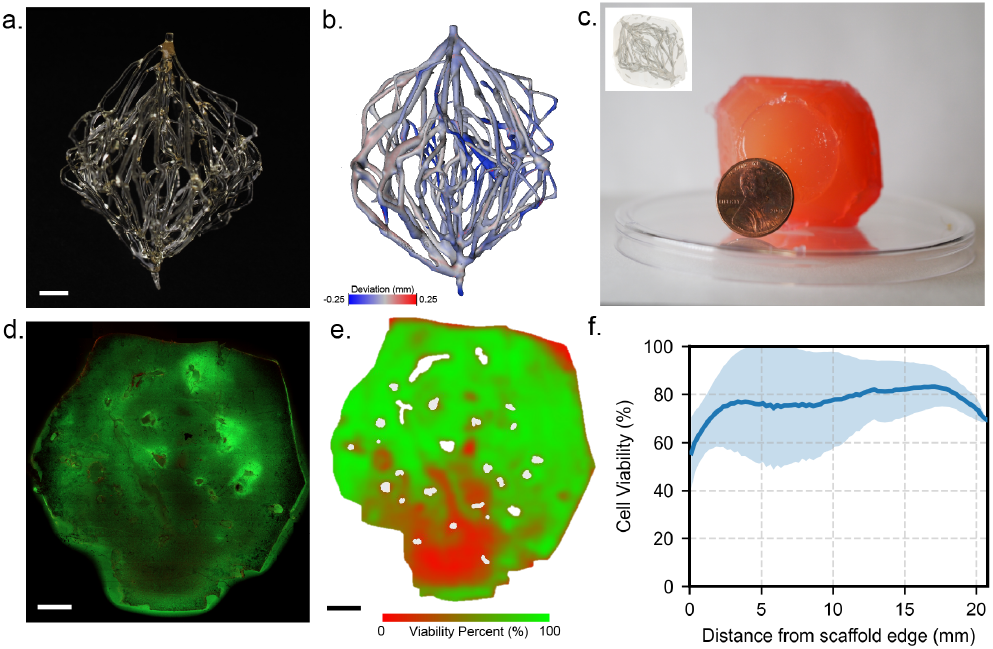
Design, fabrication, and functional validation of a biomimetic large pCAST vascular construct. **(a)** Three-dimensional rendering of a biomimetic vascular network generated using a space-colonization– based growth algorithm. Scale bar 5 mm. **(b)** Finite-element model of oxygen transport within the volumetric construct, showing predicted oxygen saturation throughout the tissue domain. **(c)** Photograph of the CLIP-printed sacrificial vascular template prior to embedding. **(d)** Representative LIVE/DEAD fluorescence image of the perfused, cell-laden construct following culture, showing spatially heterogeneous viability. Scale bar 5 mm. **(e)** Quantitative spatial viability map derived from fluorescence image segmentation. Scale bar 5 mm. **(f)** Radial viability profile plotted as a function of distance from the template edge. Mean ± SD (N=1).

Several limitations and opportunities for future work remain. The current vascular generation algorithm does not optimize local flow uniformity or shear stress, motivating future incorporation of coupled fluid– transport optimization to further improve oxygen homogeneity while maintaining physiologic shear environments throughout the network. Second, all studies here employed conventional cell culture media, which lacks the oxygen-carrying capacity of blood. Incorporating blood or blood-mimetic oxygen perfusates represents an important future direction, as hemoglobin-mediated oxygen binding and release may further extend viable tissue dimensions without increasing perfusion rates beyond physiological, an essential consideration for translating engineered vascular networks toward organ-scale perfusion and transplant-relevant environments. Looking ahead, these engineering advances also create a natural bridge to biological maturation of the channels themselves: lining pCAST-fabricated networks with endothelial cells (and supporting mural/perivascular populations) would enable barrier function, flow-responsive remodeling, and vascular signaling that more closely mimics native microcirculation. This combination of designed transport and living endothelium would position pCAST as a platform for vascularized organoids and tumor models in which hypoxia, nutrient gradients, and perfusion can be tuned with precision, enabling more faithful studies of tumor evolution, angiogenic dynamics, and transport-limited drug penetration and efficacy. In turn, such controllable, perfused microphysiological systems could strengthen drug screening and discovery by improving extrapolation from *in vitro* response to *in vivo* performance. Finally, the same framework—scalable fabrication of hierarchical perfusable networks coupled to predictive transport modeling—suggests broader applicability beyond tissue engineering, including next-generation bioreactors for biologics and cell-based manufacturing where oxygen delivery, metabolite removal, and shear remain fundamental constraints on productivity and product quality.

## Acknowledgments

Large language models were utilized to assist with conceptualization, coding optimization, manuscript revisions, and pre-peer review critiquing. All scientific content was generated and verified by the authors.

## Author contributions

Conceptualization: IAC, CAK, YLT, DIA, AN, EEH, AK, MTD, BS, MASS, JWM, ESGS, JMD

Simulation: IAC, CAK, ESGS, JMD

Experimentation: IAC, CAK, YLT, DIA, AN, EEH, AK Cell Culture: YLT, DIA, AN

Funding acquisition: MTD, BS, MASS, JWM, ESGS, JMD

Project administration: MTD, BS, JWM, ESGS, JMD

Writing – original draft: IAC, CAK, YLT

Writing – review & editing: IAC, CAK, YLT, DIA, AN, EEH, AK, MTD, BS, MASS, JWM, ESGS, JMD

## Data and materials availability

All data and materials corresponding to the research is contained within this manuscript and supplementary materials.

