## Supplementary material for "Photopatterned Sacrificial Vascular Architectures for Large Tissue-Scale Oxygenation": Methods: FinalMethods.pdf

### Materials and Methods:

#### Flow and Oxygen Transport Finite Element Analysis (FEA) Model

Flow and oxygen transport simulations were performed in COMSOL Multiphysics 6.3, utilizing the "Free and Porous Media Flow, Brinkman", "Transport of Diluted Species in Porous Media", and "Reacting Flow, Diluted Species" modules.

Momentum transport calculations were performed using the Brinkman Equations (Eq. S2-S4). The cellularized gel tissue was treated as a high-porosity, low-permeability Darcian porous medium. The perfusion channels were treated as freely flowing channels. External walls were given the no-slip boundary condition. Internal boundaries between freely flowing and porous domains were given the porous slip boundary condition.

Continuity equation for an incompressible fluid:

$$\nabla \cdot \boldsymbol{u} = 0 \qquad \text{(Eq. S1)}$$

Modified Navier-Stokes equation for incompressible flow in a Darcian porous medium:

$$\frac{\rho}{\epsilon_p}\left(\frac{\partial \boldsymbol{u}}{\partial t} + (\boldsymbol{u} \cdot \nabla)\frac{\boldsymbol{u}}{\epsilon_p}\right) = \nabla \cdot (-P\boldsymbol{I} + \boldsymbol{K}) - \frac{\mu}{\kappa}\boldsymbol{u} + \rho \boldsymbol{g} \qquad \text{(Eq. S2)}$$

Viscous stress tensor:

$$\boldsymbol{K} = \frac{\mu}{\epsilon_p}\left((\nabla \boldsymbol{u} + (\nabla \boldsymbol{u})^T) - \frac{2}{3}(\nabla \cdot \boldsymbol{u})\boldsymbol{I}\right) \qquad \text{(Eq. S3)}$$

For mass transport calculations of oxygen, we employed a typical porous advection-diffusion-reaction mass balance…

$$\epsilon_p \frac{\partial C}{\partial t} + \frac{\partial(\rho C_P)}{\partial t} - D_{eff}\nabla^2 C + \boldsymbol{u} \cdot \nabla C = -\frac{V_{max}C}{K_M + C}\mathcal{H}(C - C_{cr}) \qquad \text{(Eq. S4)}$$

where $D_{eff}$ is the effective diffusivity in porous medium, the reaction term is governed by Michaelis-Menten kinetics, and $\mathcal{H}(C - C_{cr})$ is a smoothed step function which equals zero below some critical oxygen concentration $C_{cr}$ (implemented for numerical stability and to estimate consumption reduction upon necrosis due to severe hypoxia).

#### Simulated Cell Viability Estimation

Cell viability was simply found by evaluating $\mathcal{H}(C - C_{cr})$ as a scalar metric within the 3D medium. To match experimental live/dead-stained microscopy observations showing that viability did not rise above 80%, the maximum value of this metric was capped at 0.8 for the simulations. Although a value of $0.05*C_{atm}$ was chosen (5% of the oxygen concentration present in air) as this threshold, we observed that some cells could survive conditions as low as $0.001*C_{atm}$ for prolonged periods.

#### Computational Model Parameters

All computational parameters can be found in Table S1.

#### pCAST Template Continuous Liquid Interface Production (CLIP)-Printing

*pCAST resin formulation:* 4-acryloylmorpholine (ACMO; Sigma-Aldrich) was mixed with 5% (w/v) phenylbis(2,4,6-trimethylbenzoyl)phosphine oxide (BAPO; Sigma-Aldrich), unless otherwise specified. The mixture was sonicated for ≈15 min to ensure complete dissolution of BAPO, stored in the dark, and used for printing on the same day. A trace amount of resin dye (DecorRom) can also be incorporated into the resin to tint it and enhance visualization.

*pCAST CLIP-printing:* pCAST templates were designed using Autodesk Fusion software, with slicing and support generation performed in Carbon software. All template geometries were printed on the prototype S2 high-resolution CLIP printer (Carbon), except for the large-volume biomimetic construct, which was printed on the Carbon M1 to accommodate its larger build area. After printing, excess resin in template crevices was gently removed with a pressurized air gun. The templates were then briefly washed with isopropyl alcohol (IPA; Sigma-Aldrich), dried again with the air gun, and post-cured in a UV curing oven (APM Technica) for 4 min. Templates were stored in the dark until use.

*Brightfield visualization of template and channel dimensions:* Printed templates and resulting hollow channel dimensions within bulk matrices after sacrificing the templates were imaged on the Olympus DSX1000 microscope (Evident Scientific) to assess fabrication resolution and fidelity.

**Fabrication of Perfusion Chambers**

Cellular perfusion chambers and covers were designed in Autodesk Fusion, with slicing performed using Carbon software. Parts were fabricated on a Carbon M1 printer using biocompatible KeySplint Hard resin (Keystone Industries). Gaskets providing an airtight seal between the perfusion chamber and cover were printed to size on a Carbon M2 printer using Carbon EPU 40 resin. Screws were used to compress the gasket and form a leak-tight seal between the perfusion chamber body and cover, while 20-gauge needles were press-fit into the printed components, yielding a fully enclosed chamber suitable for perfusion.

Oxygen-tracking perfusion chambers were fabricated using the same design, printing, and assembly procedures described above, with one modification. The bottom of the chamber was left open during printing, and a glass microscope slide was subsequently cured into place to form the chamber base. An oxygen-sensing foil (PreSens) was affixed to the glass surface using double-sided transparent 3M tape (3M). All remaining components, materials, and assembly steps including gasket compression via screws and press-fit integration of 20-gauge needles were identical to those used for the standard perfusion chambers.

For all cell experiments, all components were sterilized by autoclaving prior to use, with the exception of the gaskets and the oxygen-sensing perfusion chamber containing the oxygen-sensitive sensor foil, which were sterilized by immersion in 70% ethanol (LabChem) and drying before use.

**pCAST Template Dissolution Studies**

pCAST resin discs (5mm × 1mm) containing varying concentrations of BAPO (0.5–5% w/v) were printed and prepared for dissolution testing. One side of each disc was adhered to a thin glass slide using superglue to facilitate handling, and the initial mass of each template–slide assembly was recorded. The template–slide assemblies were fully immersed in deionized (DI) water at room temperature. At defined time points, templates were carefully removed, gently dried with an air jet to remove excess surface water and weighed. This process was repeated until the measured mass reached a constant value, indicating complete dissolution of the pCAST template. The mass of the glass slide and superglue was subtracted from all measurements to calculate the temporal change in template mass to determine the template dissolution profiles as a function of BAPO concentration.

**Oxygen Mapping within pCAST Constructs**

Oxygen levels within 3D hydrogel constructs were measured using the VisiSens TD oxygen imaging system (PreSens) together with oxygen sensor foil SF-RPSu4 (PreSens) (see “Fabrication of Perfusion Chambers” for chamber details). The VisiSens TD camera was positioned above the chamber to image the oxygen sensor foil, and the system was calibrated according to the manufacturer’s instructions using 0% (sodium sulfite solution) and 21% oxygen standards. Time-lapse measurements were acquired under perfusion conditions. Oxygen maps were generated with VisiSens Analyzer software, which converts luminescence lifetime signals from the sensor foil into quantitative oxygen concentrations (µM) or percent saturation. During imaging, care was taken to minimize light exposure and movement of the constructs. Measurements were conducted at room temperature for acellular experiments and at 37 °C for cellular experiments.

**Synthesis and Characterization of Gelatin Methacrylate (GelMA)**

*GelMA synthesis:* GelMA was synthesized following a previously described procedure (1). Briefly, 60 g of gelatin from cold-water fish skin (CWF gelatin; Sigma-Aldrich, G7041) was dissolved to 20% (w/v) in 300 mL of 0.1 M carbonate–bicarbonate buffer at 50 °C for ≈1 h. The pH was adjusted to 10 using 10 M sodium hydroxide (Sigma-Aldrich, 415413), and the solution was then heated to 70 °C. 5 mL of methacrylic anhydride (Sigma-Aldrich, 276685) was added with stirring at 100–200 rpm, and the reaction was allowed to proceed for 2 h at 70 °C under continuous stirring, with the reaction beaker covered by aluminum foil to minimize evaporation. After cooling to room temperature, the reaction mixture was precipitated in excess volume (≈3x volume) of absolute ethanol (≥99.5%, Sigma-Aldrich, 459844). The GelMA precipitate was collected, dried for more than 24 h, redissolved in 100 mL DI water, and reheated to approximately 80 °C for ≈1–2 h with stirring to remove residual ethanol. The GelMA solution was cooled and stored at 4 °C until use. For biological experiments, the GelMA solution was sterilized by autoclaving prior to use.

*Determination of GelMA solution concentration:* An aliquot of known volume was lyophilized overnight, and the resulting dry mass was measured to calculate the GelMA concentration. The stock GelMA solution was then diluted to the desired concentration for subsequent experiments.

*Proton Nuclear Magnetic Resonance ($^1$H-NMR) spectroscopy:* To determine the degree of methacrylation, the synthesized GelMA was characterized using $^1$H-NMR. A small aliquot of the GelMA solution was lyophilized overnight to obtain the dry polymer. Dried GelMA and native cold-water fish (CWF) gelatin (5 mg each) were dissolved separately in 1 mL $D_2O$ (Sigma-Aldrich, 151882). $^1$H-NMR spectra were acquired on a 600 MHz spectrometer (Varian). Spectra were phase- and baseline-corrected, and peak integrals were obtained using MestReNova. The aromatic amino acid signals (7.05–7.25 ppm in $D_2O$) served as an internal reference to normalize the lysine amino acid signals (2.80–2.90 ppm in $D_2O$) for both samples[2]. As methacrylic anhydride reacts with primary amines on lysine residues of gelatin to form GelMA, the degree of methacrylation reported to be 72.3 % was calculated as the percentage decrease in the normalized lysine signal in GelMA relative to CWF gelatin (Equ. S5) (2)

$$Degree\ of\ methacrylation = \left(1 - \frac{Normalized\ lysine\ integral\ signal\ of\ GelMA}{Normalized\ lysine\ integral\ of\ CWF\ gelatin}\right) \times 100\% \qquad (S5)$$

*Unconfined compression mechanical testing:* GelMA prepolymer solutions at 10–20% (w/v) containing 2 mM lithium phenyl-2,4,6-trimethylbenzoylphosphinate (LAP; Sigma-Aldrich) were prepared in 1× Dulbecco's phosphate-buffered saline (DPBS; Corning, 21-031-CV). The solutions were dispensed into cylindrical molds (10 mm diameter × 3 mm height) and photocrosslinked for 2 min under 395–405 nm irradiation using a handheld UV flashlight (UltraFire UV). The GelMA discs were demolded and equilibrated by soaking in 1× DPBS overnight at room temperature. Immediately before testing, the discs were gently dabbed dry with a Kimwipe to remove excess surface liquid. Unconfined compression testing was conducted on a mechanical tester (Instron 5565) with a 100 N load cell at a strain rate of 20% $min^{-1}$. The compressive modulus was obtained from the slope of the linear region of the stress–strain curve between 0 and 5% strain of 3.1 kPa.

**Cell Culture**

All cells were maintained at 37 °C and 5% $CO_2$ in the incubator. Immortalized Human Embryonic Kidney 293T (HEK-293T) cells and primary Human Neonatal Dermal Fibroblasts (HNDF) (gifted by Prof. Shaomin Tian, University of North Carolina at Chapel Hill and Prof. Mark Skylar-Scott, Stanford University) were cultured in Dulbecco's Modified Eagle's Medium (DMEM; Corning, 10-013-CV) supplemented with 10% Fetal Bovine Serum (FBS; Sigma-Aldrich, F2442) and 1% Antibiotic-Antimycotic (Anti-anti, Sigma-Aldrich, A5955). Cultures were maintained with media changes every 2 days and passaged at a 1:4 ratio upon confluence. Cell culture flasks were rinsed with 1× DPBS prior to detachment with 0.05% Trypsin-EDTA (Sigma-Aldrich, T4049) for 5 min at 37 °C. The reaction was neutralized with 10% FBS solution, followed by centrifugation at 1000 rpm for 4 min before resuspension and seeding.

**Fabrication of Perfusable Constructs with pCAST**

Perfusable constructs were generated using pCAST constructs as sacrificial templates. Printed pCAST templates were positioned within perfusion chambers by inserting their ends into the inlet and outlet needles. Pre-polymer solutions were then pipetted around the template and cured in situ to form the surrounding bulk matrix.

For hydrogel constructs, 10% (w/v) GelMA containing 2 mM lithium phenyl-2,4,6-trimethylbenzoylphosphinate (LAP) was introduced into the chamber and photocrosslinked for 2 min under 395–405 nm irradiation (UltraFire). Upon gelation, the pCAST template was allowed to dissolve within the aqueous GelMA matrix for 5–10 min. The inlet and outlet needles were then removed, flushed with 1× DPBS to clear residual template material, reattached, and the dissolved template was fully removed by perfusion with 1× DPBS at approximately 10 µL/min, yielding a hollow, perfusable channel network.

For elastomeric constructs, SYLGARD™ 184 (Dow) was mixed at a 10:1 base-to-curing-agent ratio and pCAST around the pCAST template within the chamber. The PDMS was cured at 85 °C overnight, after which the solidified construct was removed from the chamber, immersed in deionized water, and sonicated to dissolve the sacrificial template. Residual template material was cleared by flushing the channels with a needle and syringe.

Perfusable channels were optionally flushed with resin dye (DecorRoom) for visualization and imaging.

**Extended Perfusion of Cellularized pCAST Constructs**

Cells were harvested as described in the "Cell Culture" section, resuspended in 1× DPBS, and combined with reagents to yield a final cell-laden pre-polymer solution comprising 10% (w/v) GelMA, 2 mM LAP (sterile-filtered), and 1% (w/v) fibrinogen (bovine plasma, sterile-filtered; Sigma-Aldrich, F8630). This composition was used across all reported constructs unless otherwise noted. The cell-laden pre-polymer solution was pipetted into the chamber with the pCAST template, and immediately photocrosslinked for 2 min under 395–405 nm irradiation (UltraFire). As detailed in the section "Fabrication of Perfusable Constructs with pCAST" for aqueous constructs, the pCAST template was dissolved and removed by low-rate DPBS perfusion, yielding a hollow, perfusable channel. Additionally, to clear any residual template from the channels, the photocured constructs were flushed thoroughly for >5 min with 1× DPBS at 150–250 µL/min via an external peristaltic pump (Golander BT100S-1) prior to connection to the main perfusion circuit with cell culture medium.

For perfused samples, cellularized constructs were enclosed under a gasketed cover to maintain sterility, reduce evaporation and limit oxygen diffusion across the chamber surfaces, leaving the perfusion channels as the primary exchange pathway. The enclosed perfusion chambers were connected via gas-permeable silicone tubing (1 mm ID; Golander, 1×0.92-S) to an external peristaltic pump and a medium reservoir fitted with a sterile filter port for gas exchange. Unless otherwise specified, constructs were perfused at 250 µL/min for 5 days using the medium appropriate for each cell type (see "Cell Culture"). The perfusion circuit was maintained at 37 °C and 5% $CO_2$ within the cell culture incubator. For each 40 mL medium reservoir, 1–3 chambers were connected, and complete medium changes were performed every 2 days.

For non-perfused controls lacking channels, cellularized constructs of identical composition were removed from the chamber immediately after photocuring and immersed in static culture medium in a 12-well plate. Controls were incubated for 5 days with complete medium exchanges every 2 days.
After 5 days of perfusion, constructs were sliced into approximately 1–2 mm-thick cross-sections, then stained for cell viability and imaged as described in "Cell Viability Staining, Imaging, and Postprocessing".

**Cell Viability Staining, Imaging and Postprocessing**

*LIVE/DEAD Staining:* Cell viability was assessed using LIVE/DEAD staining (Thermo Fisher Scientific, L3224). Cellularized samples were briefly rinsed with 1× DPBS and incubated at 37 °C for 20 min in 1× DPBS containing calcein AM (0.5 μL/mL) and ethidium homodimer-1 (EthD-1; 2 μL/mL) to label live and dead cells, respectively.

*Microscopy Imaging:* LIVE/DEAD imaging on the cross sections of the perfused cellularized constructs was done on the Zeiss LSM980 laser scanning microscope. Confocal Z-stacks were acquired using 488 nm excitation with a 1 Airy unit pinhole. Volumes spanning 300 μm were collected with a 58.7 μm Z-step (6 slices), centered on the region of interest, with detector settings held constant across samples. LIVE/DEAD fluorescence imaging to assess pCAST template biocompatibility, matrix stiffness effects, and endothelialization studies was conducted using a Leica THUNDER fluorescence microscope; acquisition parameters are detailed in the corresponding sections.

*LIVE/DEAD Image Postprocessing:* Unless otherwise noted, maximum-intensity projections of Z-stacks were used for analysis. Images were processed in ImageJ using a custom macro to segment and count calcein AM–positive (live) and EthD-1–positive (dead) cells. Cell viability was calculated as the ratio (in %) of live cells to all cells.

*Cell Viability Heatmap Generation:* Spatial viability maps were generated using a custom Python-based image analysis pipeline developed in-house. Confocal image stacks were first preprocessed to remove background and normalize intensity across fields of view. Live and dead fluorescence channels were segmented using fixed intensity thresholds applied consistently across all samples. Binary masks were used to compute local viability as the fraction of live signal relative to total (LIVE/DEAD) signal. Viability values were then spatially averaged over a defined grid to generate continuous two-dimensional heatmaps representing cell survival within the construct. To enable quantitative comparison across conditions, pixel-to-distance calibration was applied using known imaging scale factors, and heatmaps were smoothed using a Gaussian filter to reduce pixel-level noise while preserving mesoscale gradients. Resulting viability heatmaps were used to extract distance-dependent viability profiles relative to perfused channels and to directly compare experimental viability patterns with FEM-predicted oxygen distributions.

**Statistical Analysis**

Unless otherwise noted, all experiments were conducted with at least N = 3 independent samples per condition. Statistical analyses were performed using OriginLab software. Unlesss otherwise stated, one-way ANOVA followed by Tukey's post-hoc test was applied to assess statistical significance. Statistical significance is indicated as follows: n.s. (not significant), *$P < 0.05$, **$P < 0.01$, ***$P < 0.001$, and ****$P < 0.0001$.
