## Supplementary Figures for "Photopatterned Sacrificial Vascular Architectures for Large Tissue-Scale Oxygenation": Final Supplementary Figures.pdf

**Supplementary Figures and Tables**

**Table S1. Model parameters for oxygen transport in perfused pCAST constructs.** Parameters defining inlet oxygen concentration, hydrogel properties, perfusion conditions, and Michaelis–Menten cellular oxygen consumption used in FEM simulations and experimental comparisons. Temperature-dependent constants were adjusted for oxygen mapping and cell viability experiments as indicated.

| Parameter | Value | Description | Reference |
|---|---|---|---|
| $C_{inlet}$ | 0.21 mol/m$^3$ | Perfusion channel inlet oxygen concentration | Determined experimentally |
| $k°_H$ | $1.3 \times 10^{-8}$ m·s$^2$·mol/kg$^2$ | Henry's Law constant at 22°C | 1 |
| T | 295.15 K for $O_2$ mapping 310.15 K for cell viability mapping | System temperature | By design |
| $d(\ln(k_H))/d(1/T)$ | 1500 K | Temperature dependence constant for Henry's Law exponential term | 1 |
| $MW_{O2}$ | 32 g/mol | $O_2$ molecular weight | |
| $\varepsilon_p$ | 0.9 | Hydrogel porosity | Estimated from water mass % |
| $\kappa$ | $10^{-14} – 10^{-17}$ m$^2$ | Hydrogel hydraulic permeability | 2 |
| $D_{O2,water}$ | $2.0–2.8 \times 10^{-9}$ m$^2$/s depending on temperature | Oxygen diffusivity in the liquid phase | 3-6 |
| $\rho_{water}$ | 1000 kg/m$^3$ | Density of water | |
| $V_{max}$ | (Cell Density) × $3.85 \times 10^{-17}$ mol/s | HEK cell maximum oxygen consumption rate | 7-10 |
| $K_M$ | $4.37 \times 10^{-2}$ mol/m$^3$ | HEK cell Michaelis-Menten constant | 11, 12 |
| Cell Density | $15–100 \times 10^{12}$ cells/m$^3$ | Density of HEK cells in the hydrogel | By design |
| $C_{cr}$ | $11.2 \times 10^{-3}$ mol/m$^3$ (~5% of atmospheric oxygen) | Critical oxygen concentration below which HEK cells experience hypoxia | 13-15 |
| Q | $2.5 \times 10^{-10} – 4.17 \times 10^{-9}$ m³/s | Inlet flow rate | By design |
| $W_{channel}$ | $4 \times 10^{-4}$ m | Perfusion channel thickness | By design |
| $d_{chan2sens}$ | $2.5 \times 10^{-4}$ m | Distance between the perfusion channels to the oxygen-sensing surface | By design |

**Table S2. Fabrication times of pCAST scaffolds printed using CLIP-based vat photopolymerization (VPP).** With VPP, print duration primarily scales with scaffold height and cross-sectional area, and is independent of geometric complexity or density. “Max. height” denotes the total height of the scaffold in its printing orientation, accounting for channel inlets/outlets and all required support structures. “No. of copies per print” indicates the number of scaffold replicates that can be accommodated on the printer build area.

| pCAST Geometry | | Channel dimensions (mm) | Max Height (mm) | Total fabrication time (min) | No. of copies per print | Fabrication time per copy (min) |
|---|---|---|---|---|---|---|
| 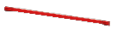 | Axial | 0.25 x 0.30 | 0.38 | 1.2 | 7 | 0.2 |
| 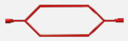 | Dual | 0.20 x 0.30 | 0.38 | 1.2 | 2 | 0.6 |
| 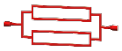 | Quad | 0.20 x 0.22 | 0.42 | 1.3 | 2 | 0.7 |
| 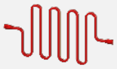 | Serpentine | 0.20 x 0.30 | 0.42 | 1.3 | 1 | 1.3 |
| 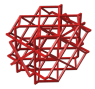 | 3D lattice | 0.18Ø | 10.50 | 27.2 | 3 | 9.1 |
| 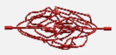 | 3D biomimetic | 0.19Ø | 4.20 | 9.8 | 1 | 9.8 |
| 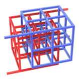 | Interlocked cubes | 0.24 x 0.24 | 7.00 | 11.4 | 1 | 11.4 |
| 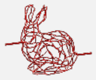 | Stanford bunny | 0.40Ø | 10.60 | 12.9 | 1 | 12.9 |
| 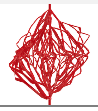 | Large-volume biomimetic | >0.25 | 37.91 | 66 | 8 | 8.25 |

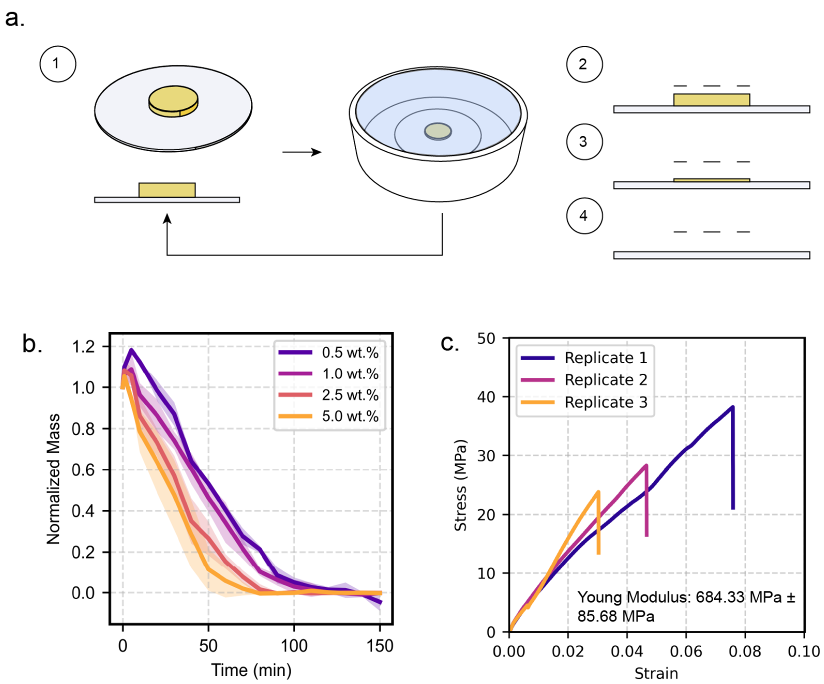


**Fig. S1. Optimization and characterization of pCAST scaffold formulation. (a)** Printability of pCAST scaffolds as a function of BAPO photoinitiator concentration. **(b)** Dissolution kinetics of pCAST scaffolds at varying BAPO photoinitiator concentrations. **(c)** Tensile mechanical characterization of pCAST scaffold.

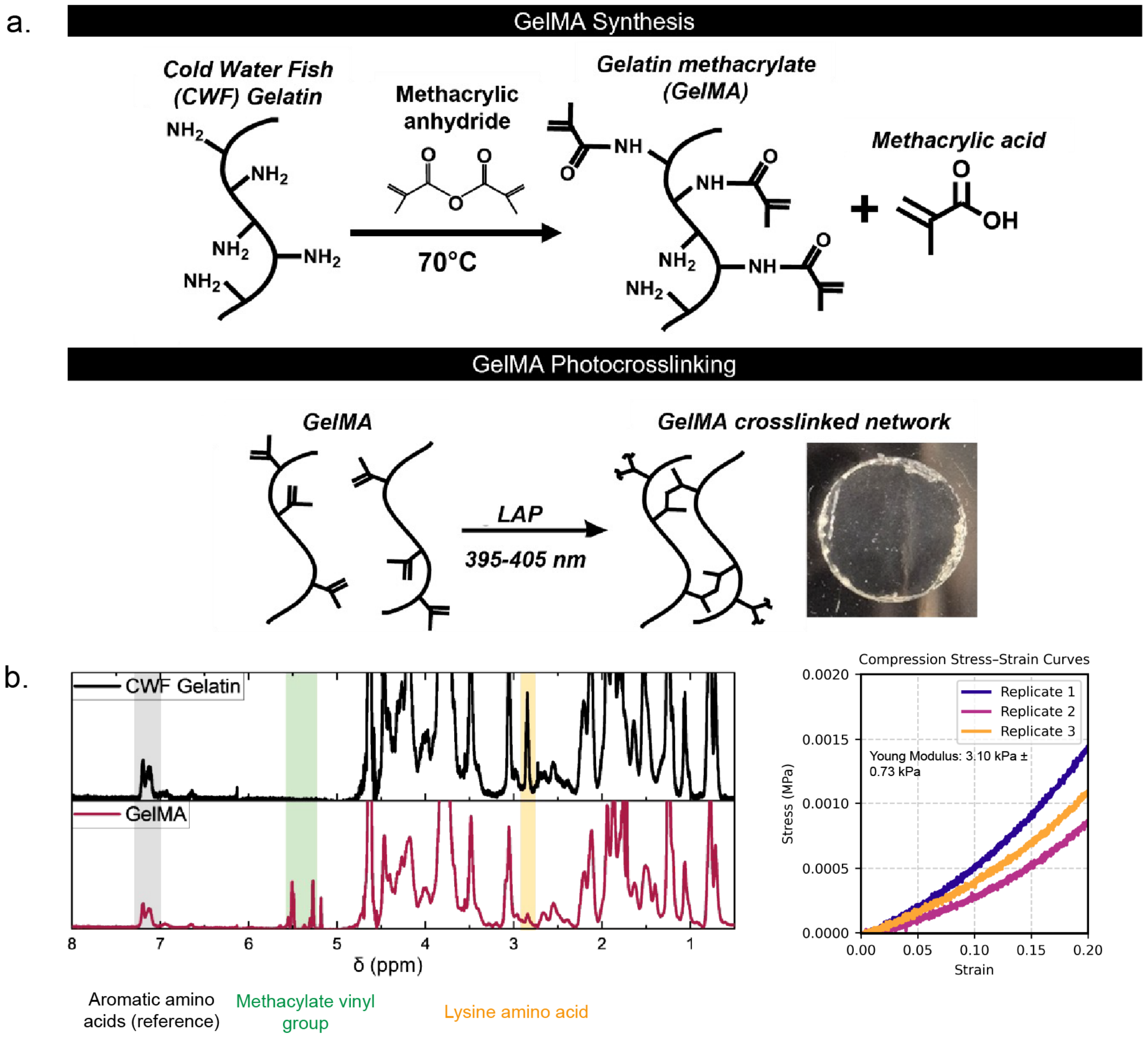


**Fig. S2. Synthesis and characterization of GelMA. (a)** Reaction schematic of GelMA synthesis and subsequent photocrosslinking. **(b)** $^1$H-NMR spectra of CWF gelatin and GelMA. **(c)** Stress–strain curves from unconfined compression testing of GelMA hydrogels at 10% (w/v).

**Table S3.** Degree of methacrylation (mean ± SD, N = 3 independent synthesis batches) and compressive modulus of GelMA hydrogels (mean ± SD, N =3).

| Degree of methacrylation | GelMA Concentration (w/v) | Compressive Modulus (kPa) |
|---|---|---|
| 72.3 ± 5.0 % | 10% | 3. 1 ± 0.73 |

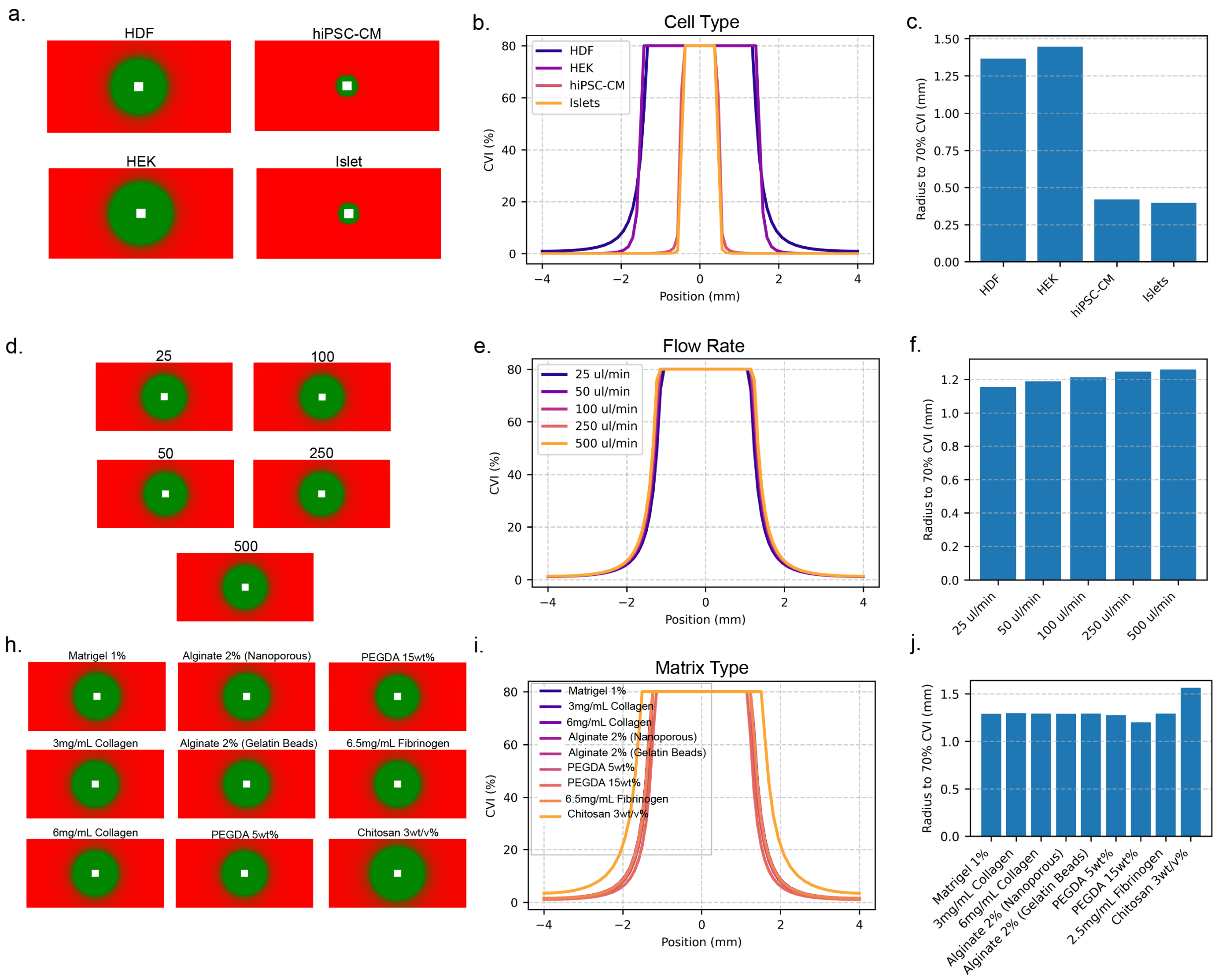


**Fig. S3. Finite-element analysis of parameters governing oxygen penetration and viability radius in perfused pCAST constructs. (a)** Simulated cross-sectional cell viability fields for different cell types (HDF, HEK-293T, hiPSC-derived cardiomyocytes, and islets) under identical vascular geometry and perfusion conditions, illustrating cell-type–dependent oxygen consumption. **(b)** Corresponding radial viability index (CVI) profiles as a function of distance from the perfused channel for each cell type. **(c)** Quantification of the radius at which CVI falls below seventy percent for each cell type. **(d)** Simulated cell viability fields for increasing perfusion flow rates (25–500 µL min$^{-1}$) at fixed cell type and geometry. **(e)** Radial CVI profiles as a function of flow rate. **(f)** Radius to seventy percent CVI as a function of perfusion rate. **(g)** Simulated cell viability fields for constructs composed of different matrix materials and formulations. **(h)** Radial CVI profiles comparing matrix-dependent oxygen transport behavior. **(i)** Quantification of the radius to seventy percent CVI across matrix types. Together, these simulations demonstrate how cellular metabolic demand, perfusion rate, and matrix properties independently and jointly regulate oxygen penetration and tissue viability in pCAST constructs.

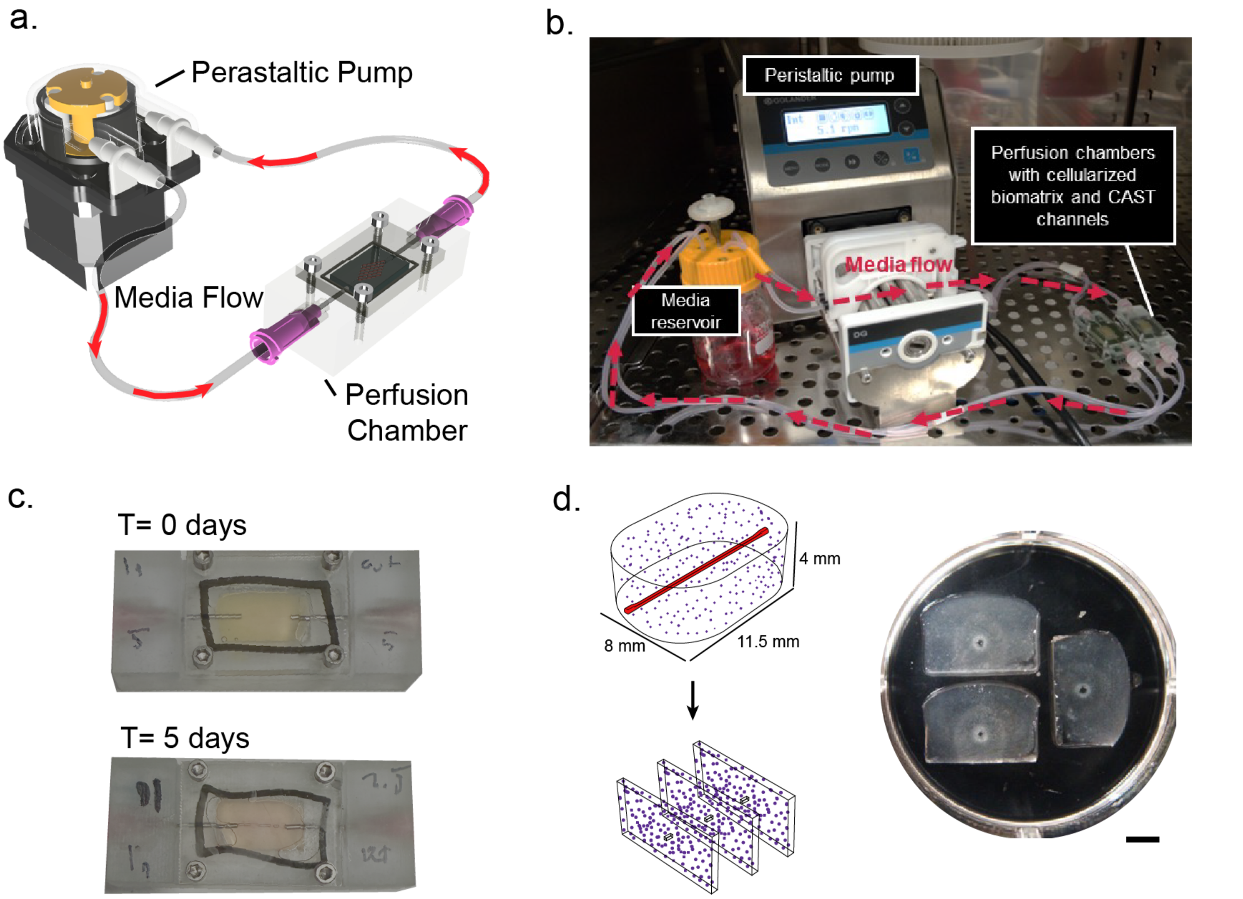


**Fig. S4. Extended perfusion setup for cell-embedded pCAST constructs. (A)** Perfusion chamber components (base, gasket, cover, inlet/outlet needle fittings) with a representative cell-laden bulk GelMA hydrogel construct. **(B)** Assembled peristaltic perfusion circuit inside a cell-culture incubator. **(C)** Representative images of cell-laden constructs at Day 0 and Day 5 under continuous perfusion. **(D)** Cross-section of representative perfused construct on Day 5 showing the pCAST-fabricated channel in the center and surrounding matrix.

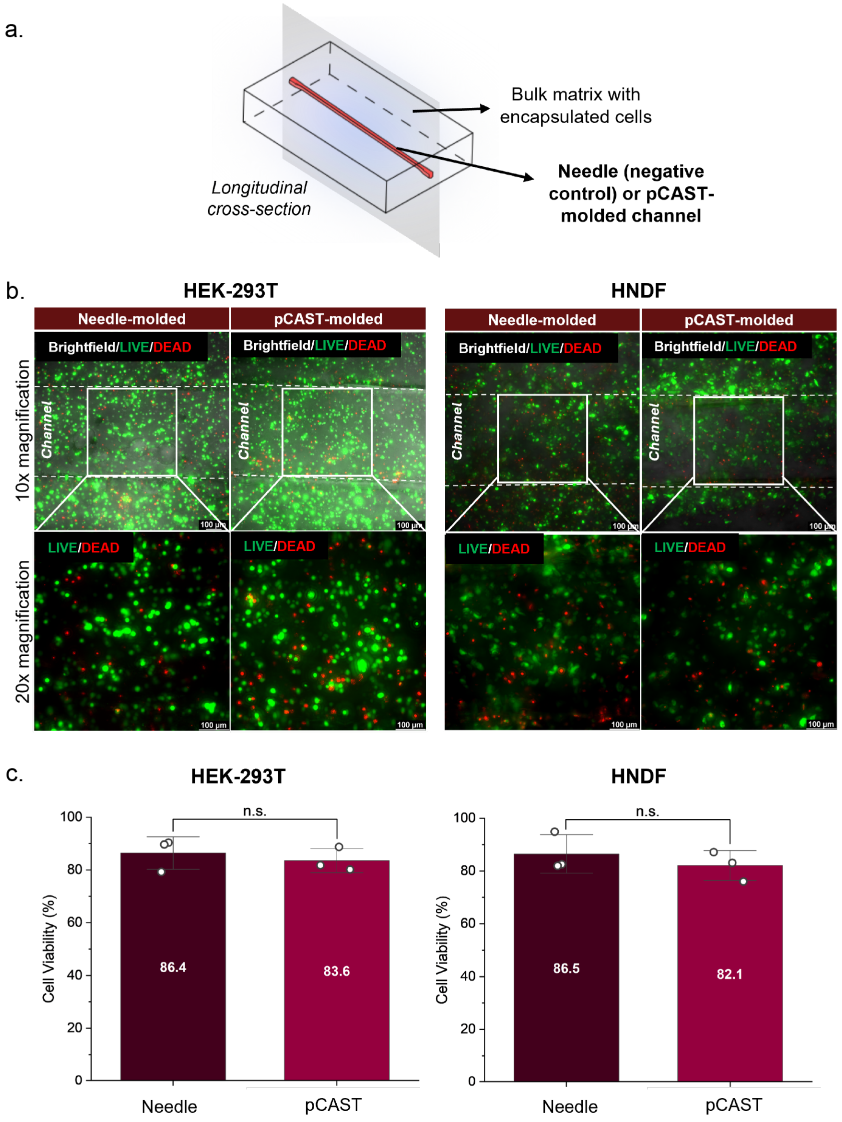


**Fig. S5. Biocompatibility of pCAST sacrificial scaffolds. (a)** LIVE/DEAD fluorescence of longitudinally bisected ("half-pipe") cellularized constructs (HEK-293T 5 × $10^6$ cells/mL; 10% w/v GelMA; 1% w/v fibrinogen; 2 mM LAP in 1× DPBS) with channels formed by either needle molding (biocompatible positive control) or a pCAST sacrificial axial scaffold, imaged after 24 h incubation in static culture media. **(b)** Quantification of cell viability at the channel wall (images taken at 20xmagnification) (mean ± SD); one-way ANOVA with Tukey's post hoc test; N = 3.

**Table S4. Literature values for perfusable cellularized construct performance.** Summary of published engineered vascular tissues with reported construct volume and minimum channel diameter. Only studies reporting both cellularized, perfusable constructs with identifiable construct dimensions were included. Construct volumes were calculated from reported width, height, and thickness dimensions when not directly provided. Values represent approximations based on reported dimensional parameters. Asterisks (*) denote minimum channel diameters determined via ImageJ analysis of published cross-sectional channel images when values were not explicitly reported.

| Label | Publication Year | Construct Volume ($cm^3$) | Min. Channel Diameter (μm) | Ref. |
|---|---|---|---|---|
| This Work | 2026 | 36 | 50-200 | n/a |
| A | 2020 | 15.5 | 300 | 14 |
| B | 2016 | 10 | 300* | 15 |
| C | 2025 | 4.75 | 400 | 16 |
| D | 2019 | 3 | 300 | 17 |
| E | 2019 | 2.5 | 400 | 18 |
| F | 2025 | 1.5 | 250 | 19 |
| G | 2025 | 1 | 1400 | 20 |
| H | 2023 | 0.67 | 250 | 21 |
| I | 2019 | 0.64 | 200* | 22 |
| K | 2024 | 0.06 | 125 | 23 |
| L | 2016 | 0.031 | 50 | 24 |
| M | 2017 | 0.012 | 50 | 25 |
| N | 2023 | 0.01 | 10 | 26 |

18. Skylar-Scott, M. A. *et al.* Biomanufacturing of organ-specific tissues with high cellular density and embedded vascular channels. *Sci. Adv.* **5**, eaaw2459.

19. Shiwarski, D. J. *et al.* 3D bioprinting of collagen-based high-resolution internally perfusable scaffolds for engineering fully biologic tissue systems. *Sci. Adv.* **11**, eadu5905.

20. Hudson, A., Shiwarski, D., Kramer, A. & Feinberg, A. Enhancing Viability in Static and Perfused 3D Tissue Constructs Using Sacrificial Gelatin Microparticles. *ACS Biomater. Sci. Eng.* **11**, (2025).

21. You, S. *et al.* High cell density and high-resolution 3D bioprinting for fabricating vascularized tissues. *Sci. Adv.* **9**, eade7923.
